# Concurrent Joint Contact in Anterior Cruciate Ligament Injury induces cartilage micro-injury and subchondral bone sclerosis, resulting in knee osteoarthritis

**DOI:** 10.1101/2024.05.08.593114

**Authors:** Kei Takahata, Yu-Yang Lin, Benjamin Osipov, Kohei Arakawa, Saaya Enomoto, Blaine A. Christiansen, Takanori Kokubun

**Author notes:** **Co-Correspondence:** ^3^Blaine A. Christiansen, University of California Davis Health, Department of Orthopaedic Surgery, Lawrence J. Ellison Musculoskeletal Research Center, 2700 Stockton Ave, Suite 2301 Sacramento, CA 95817, USA.; ^1,6^Takanori Kokubun, Ph.D., 820 San-nomiya, Koshigaya, Saitama, 343-8540. Japan.

## Abstract

**Objective:** Anterior Cruciate Ligament (ACL) injury initiates post-traumatic osteoarthritis (PTOA) via two distinct processes: initial direct contact injury of the cartilage surface during ACL injury, and secondary joint instability due to the ACL deficiency. Using the well-established Compression-induced ACL rupture method (ACL-R) and a novel Non-Compression ACL-R model, we aimed to reveal the individual effects of cartilage compression and joint instability on PTOA progression after ACL injury in mice.

**Design:** Twelve-week-old C57BL/6J male were randomly divided to three experimental groups: Compression ACL-R, Non-Compression ACL-R, and Intact. Following ACL injury, we performed joint laxity testing and microscopic analysis of the articular cartilage surface at 0 days, in vivo optical imaging of matrix-metalloproteinase (MMP) activity at 3 and 7 days, and histological and microCT analysis at 0, 7, 14, and 28 days.

**Results:** The Compression ACL-R group exhibited a significant increase of cartilage roughness immediately after injury compared with the Non-Compression group. At 7 days, the Compression group exhibited increased MMP-induced fluorescence intensity and MMP-13 positive cell ratio of chondrocytes. Moreover, histological cartilage degeneration was observable in the Compression group at the same time point. Sclerosis of tibial subchondral bone in the Compression group was more significantly developed than in the Non-Compression group at 28 days.

**Conclusions:** Both Compression and Non-Compression ACL injury initiated PTOA progression due to joint instability. However, joint contact during ACL rupture also caused initial micro-damage on the cartilage surface and initiated early MMP activity, which could accelerate PTOA progression compared to ACL injury without concurrent joint contact.

## INTRODUCTION

Osteoarthritis (OA) is a major chronic musculoskeletal disease that progresses irreversibly and negatively influences the quality of life of OA patients. OA can be divided into two types depending on the mechanisms of initiation: primary OA, which is idiopathic and generally occurs during aging, and secondary post-traumatic OA (PTOA), which is initiated by a traumatic joint injury such as a meniscus tear or ligament injury. PTOA accounts for approximately 12% of symptomatic OA patients ^1^ and has an increasing incidence rate due to the increasing popularity of high-impact sports ^2^. Therefore, elucidating the mechanisms of PTOA initiation and progression is necessary to find therapeutic strategies to prevent or slow the progression of OA.

Anterior Cruciate Ligament (ACL) injury is a known initiator of PTOA development. The annual incidence of ACL injury in the general population is 68.6 per 100,000 people ^3^, and the incidence of PTOA following ACL injury is as high as 87% ^4^. In the etiology of OA following ACL injury, concomitant injuries of other joint structures along with ACL rupture and increased mechanical stress caused by joint instability are likely contributing factors. Almost half of ACL injury patients suffer from initial damage of articular cartilage ^5^, and concomitant cartilage injury may increase OA risk up to 2.4 times at 19 years ^6^. In addition, 80-90% of patients with an acute ACL injury also show signs of subchondral bone lesions measured using magnetic resonance imaging. ^4^ Subchondral bone changes can affect cartilage degeneration via crosstalk in molecular and biomechanical interactions. ^7^.

Because the ACL restricts the anterior-posterior (AP) translation of the tibia relative to the femur ^8^, joint instability after ACL injury can induce abnormal mechanical stress, which may stimulate knee joint tissues to produce catabolic enzymes. Since knee PTOA develops several decades after ACL injury, it was assumed that secondary joint instability is the most important factor and surgical reconstruction of the ACL has been widely used to reduce the risk of PTOA development ^9^. However, a recent systematic review reported that reconstruction surgery may not affect PTOA progression ^10^. This indicates that concurrent joint contact when the ACL is injured may also play a significant role in the pathogenesis of PTOA. Acute damage to the articular chondral surface and subchondral bone could be the initiation of PTOA, however, it remains unclear which tissue would be more involved in the OA onset.

ACL transection (ACL-T) is a common model for inducing knee OA in small animals, but this method also induces unnecessary acute inflammation in articular tissues due to invasive surgical procedures ^11–14^. In contrast, the Non-Invasive Compression ACL Rupture (ACL-R) model was developed and used to explore mechanisms of PTOA progression following ACL ^15–19^. ACL-R is induced by applying compressive force to the joint, which causes anterior tibial dislocation and ACL injury. This method largely imitates acute ACL injury and knee PTOA progression after injury. However, it is difficult to separate the effects of initial overload of the articular cartilage and the secondary effects of joint instability. We recently established a novel Non-Compression ACL-R model, which is made without compression force on the cartilage surface and induces non-articular injuries other than ACL rupture ^20^. Based on the findings of this model, we hypothesized that comparing the Compression ACL-R model and the novel Non-Compression ACL-R model will allow us to investigate the individual effects of initial cartilage damage due to compression vs. secondary instability on PTOA progression.

## MATERIALS AND METHODS

### Animals and Experimental Design

Twelve-week-old C57BL/6J male mice were randomly divided into three experimental groups: Compression ACL-R (n = 49), Non-Compression ACL-R (n = 48), and Intact (n=21). Mice were euthanized at 0, 3, 7, 14, and 28 days after injury, and knee joints were dissected for analysis (Fig. 1A). In the joint laxity test, microscopic, morphological, and histological analysis at day 0 after injury, contralateral knee joints from the Non-Compression group were used as the Intact group. Mice were allowed to acclimate for 1-2 weeks in the vivarium before the start of the experiment and were cared for in accordance with the guidelines set by the National Institutes of Health (NIH) on the care and use of laboratory animals. All procedures were conducted under approved Institutional Animal Care and Use Committee protocols at Saitama Prefectural University and the University of California Davis.

**Figure 1.**
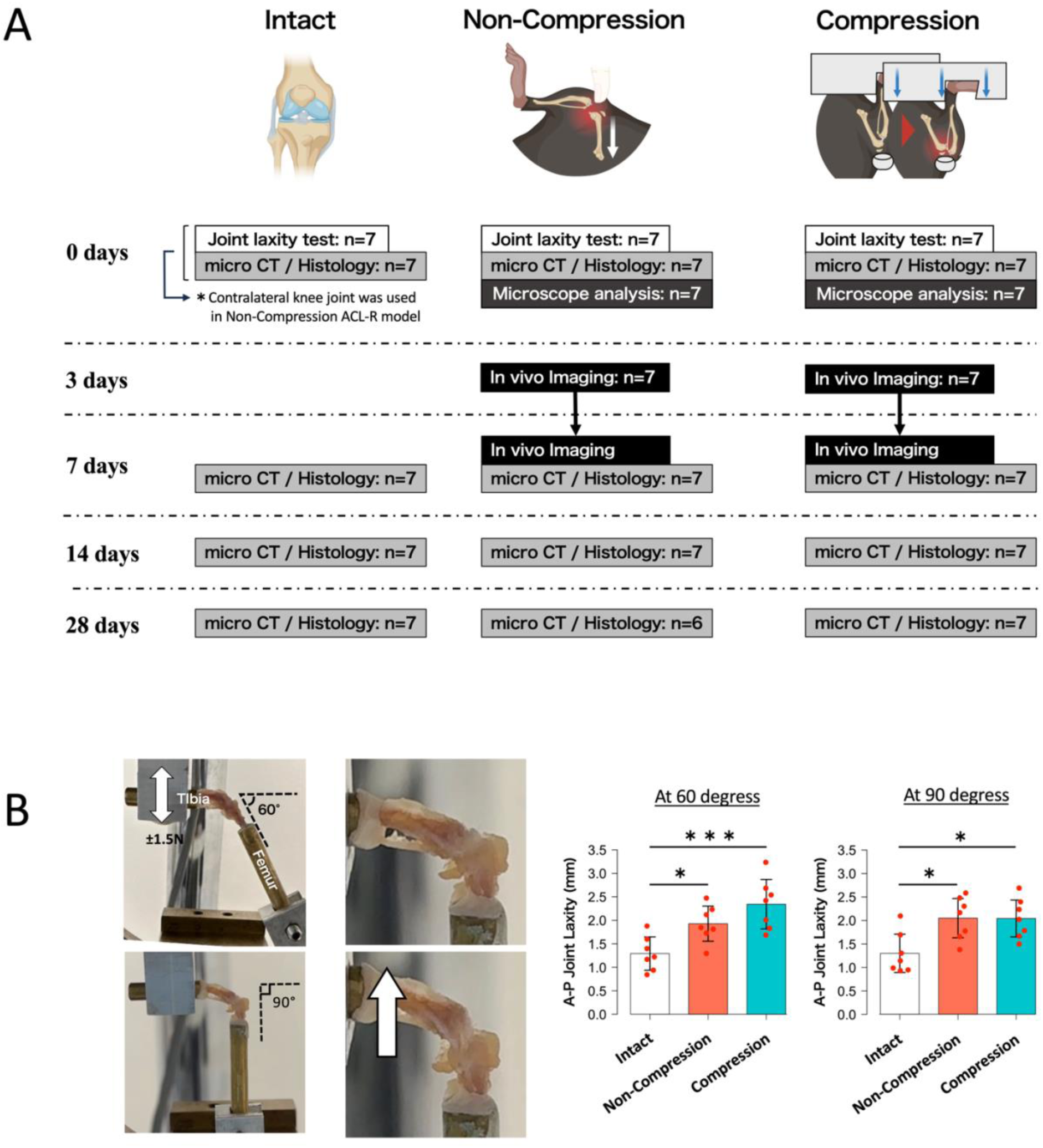
(A) Experimental design. We made the Compression ACL-R, Non-Compression ACL-R, and Intact groups. Mice were euthanized at 0, 3, 7, 14, and 28 days after injury and following analyses were performed; joint laxity test, microscopic, microCT, and histological analysis. At day 0 after injury, contralateral knee joints from the Non-Compression group were used as the Intact group. (B) A result of joint laxity test. The Compression and Non-Compression ACL-R groups significantly increased Anterior-Posterior joint laxity at both 60 ° and 90 °. However, no significant differences were observed between both injury groups. Data are presented as the mean ± 95% CI. *P< 0.05; ***P< 0.001.

### Creating Compression ACL-R and Non-Compression ACL-R Models

All procedures were performed on the right knee joint of each anesthetized mouse under 1-4% inhaled isoflurane. The Compression ACL-R model was created via tibial compression overload as previously described ^17, 21^. Right knee joints were positioned in an electromagnetic materials machine (ElectroForce 3200, TA Instruments, New Castle, DE), and a single tibial compressive overload was applied at 1 mm/s to induce ACL rupture. The Non-Compression ACL-R model was created based on our previous study ^22^. The knee joint was fixed at 90 degrees using surgical tape on a stand, and a force was slowly applied by the thumb tip of the operator along the long axis of the femur. The applied force was stopped quickly after hearing a distinct popping sound, which indicated ACL rupture. For both models, buprenorphine analgesia (0.1 mg/kg) was injected immediately following injury for pain relief.

### Anterior-Posterior Joint Laxity Testing

Knee joints were collected immediately post-injury, and femurs and tibias were embedded in brass tubes using polymethylmethacrylate as previously described ^21, 23^. Tibias were fixed to the load cell; then femurs were secured so that the angle of the knee joint was either 60 ° or 90 °. Five anterior-posterior (AP) loading cycles were applied perpendicular to the longitudinal axis of the tibia to a target force of ±1.5 N at a loading rate of 0.5 mm/s. The degree of AP joint laxity was quantified based on the difference between displacement at +0.8 N and −0.8 N.

### Microscopic Analysis of Cartilage Surface Roughness

For samples collected 0 days following injury, soft tissue was removed carefully to expose the tibial plateau, then 3D optical profilograph images of the whole tibial plateau were taken (VR-6000; Keyence, JPN). The region of interest (ROI) was set as an ellipse with 0.6-1.2 mm diameters in the center area of the medial and lateral tibial plateau (Figure 2A). The arithmetical mean height and maximum height of the cartilage surface were measured by normalizing to the contralateral intact knee of each mouse. Methods are further described in the Supplementary Materials.

**Figure 2.**
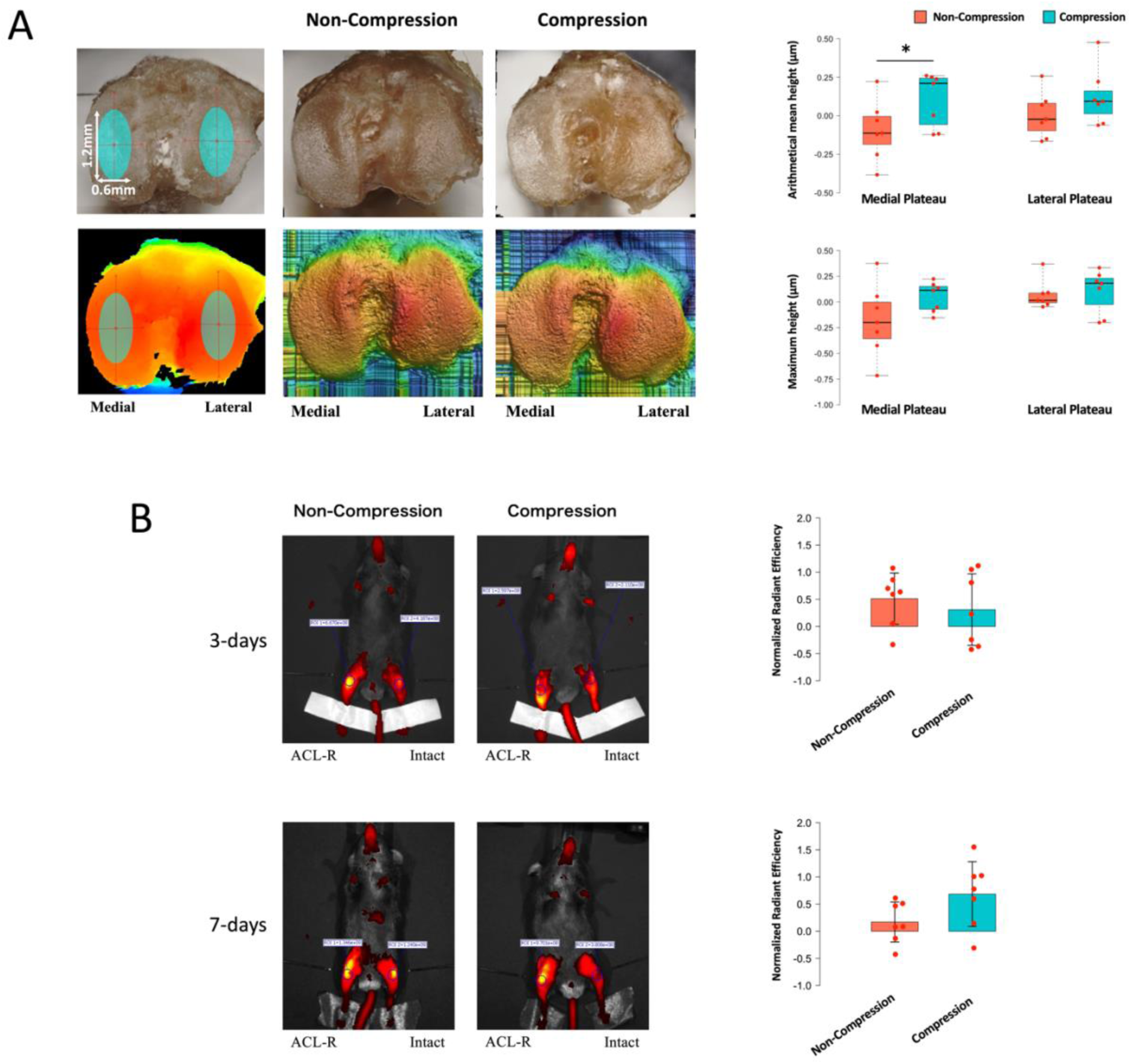
(A) Microscope Analysis of Cartilage Roughness in the Tibial Plateau at day 0 after injury. The ROI was set as an ellipse with 0.6-1.2 mm diameters in the center area of the medial and lateral tibial plateau. Compression group significantly increased the normalized arithmetical mean height in the medial compartment compared with the Non-Compression group. (B) Quantification of MMP Fluorescence Intensity. Normalized MMP-induced fluorescence intensity around ACL-R knees increased in the Compression, especially at 7 days following injury. However, there were no significant differences between these groups. Data are presented as the median ± interquartile range. *P< 0.05.

### Fluorescent Reflectance Imaging (FRI)

At 3 and 7 days post-injury, mice were imaged in vivo using an optical imaging system (IVIS Spectrum, PerkinElmer, Waltham, MA), after IV administration of a near-infrared probe that is activated by matrix-metalloproteinases (MMPSense 680, PerkinElmer, Waltham, MA). The ROI was set as a circle of 0.7 mm^2^ that surrounded the tibial tuberosity to the superior border of the patella, and image processing and quantification were performed via IVIS Living Image software as previously described ^24^. To unify the mouse-to-mouse variation in the delivery of the fluorescent probe, the radiant efficiency of the ACL-R knee was normalized to the contralateral intact knee of each mouse. Methods are further described in the Supplementary Materials.

### Micro-Computed Tomography Analysis of Osteophyte Formation and Epiphyseal Bone Microstructure

For analysis of osteophyte formation at 7, 14, and 28 days after injury, ROI contouring included all heterotopic mineralized tissue around the joint, as well as the patella, fabellae, and menisci. The average bone volume in the Intact group was used as the “baseline” volume to calculate osteophyte formation in the Compression and Non-Compression groups.

We also assessed the morphological changes in epiphyseal trabecular bone at 0, 7, 14, and 28 days after injury. The ROI was designed as the femoral and tibial trabecular bone enclosed by the growth plate and subchondral cortical bone plate (Figure 3B). Additionally, whole subchondral bone including the cortical bone plate and trabecular bone in the medial and lateral tibial compartments were also measured (Figure 4A). We quantified apparent bone mineral density (BMD, g/cm^3^), bone volume per total volume (BV/TV, %), trabecular number (Tb.N, 1/mm), trabecular thickness (Tb.Th, mm), and trabecular separation (Tb.Sp, mm).

**Figure 3.**
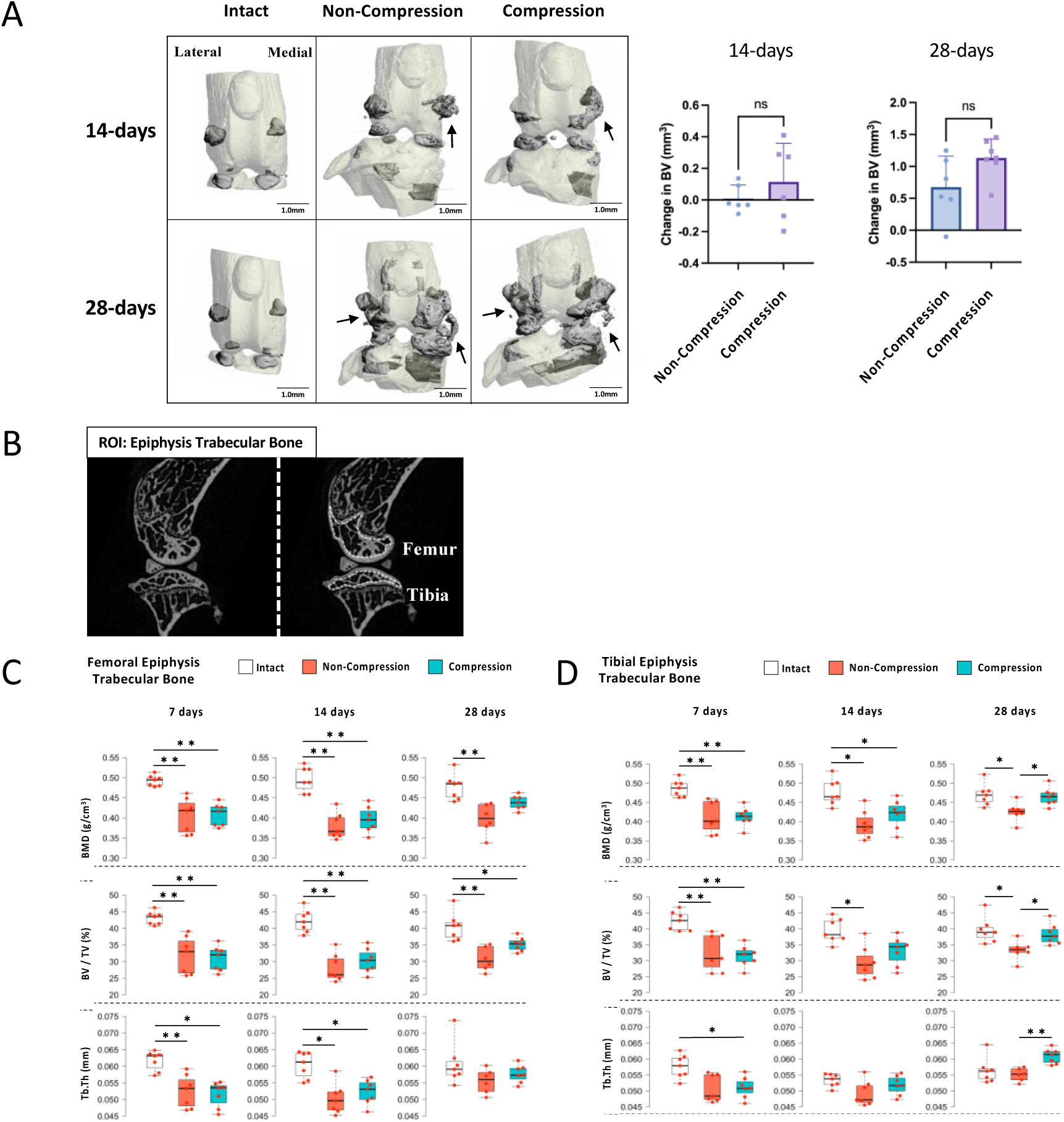
(A) Reconstructed images obtained by microCT and measurement of osteophyte formation. Only a small amount of mineralized osteophyte at 14 days and a greater volume of ectopic mineralized tissue in the medial femoral condyle and tibial plateau of injured joints at 28 days were observed in the Compression and Non-Compression groups. However, there were no significant differences. Black arrows show the osteophyte. (B) The ROI to measure the tibial and femoral epiphysis trabecular bone and (C) the result of microCT Analysis. Femoral epiphysis trabecular bone significantly decreased BMD, BV/TV, and Tb.Th in the Compression and Non-Compression groups compared with the Intact group at 7 days. At 28 days, no significant difference in the BMD was observed between the Intact and the Compression groups. However, the Non-Compression group was still significantly lower than the Intact group at the same timepoint. Tibial epiphysis trabecular bone also showed a significant decrease in BMD and BV/TV in both injury groups at 7 days. However, at 28 days, the BMD and BV/TV in the Intact and Compression groups were significantly greater than in the Non-Compression group. Data are presented as the median ± interquartile range. *P< 0.05. **P< 0.01.

**Figure 4.**
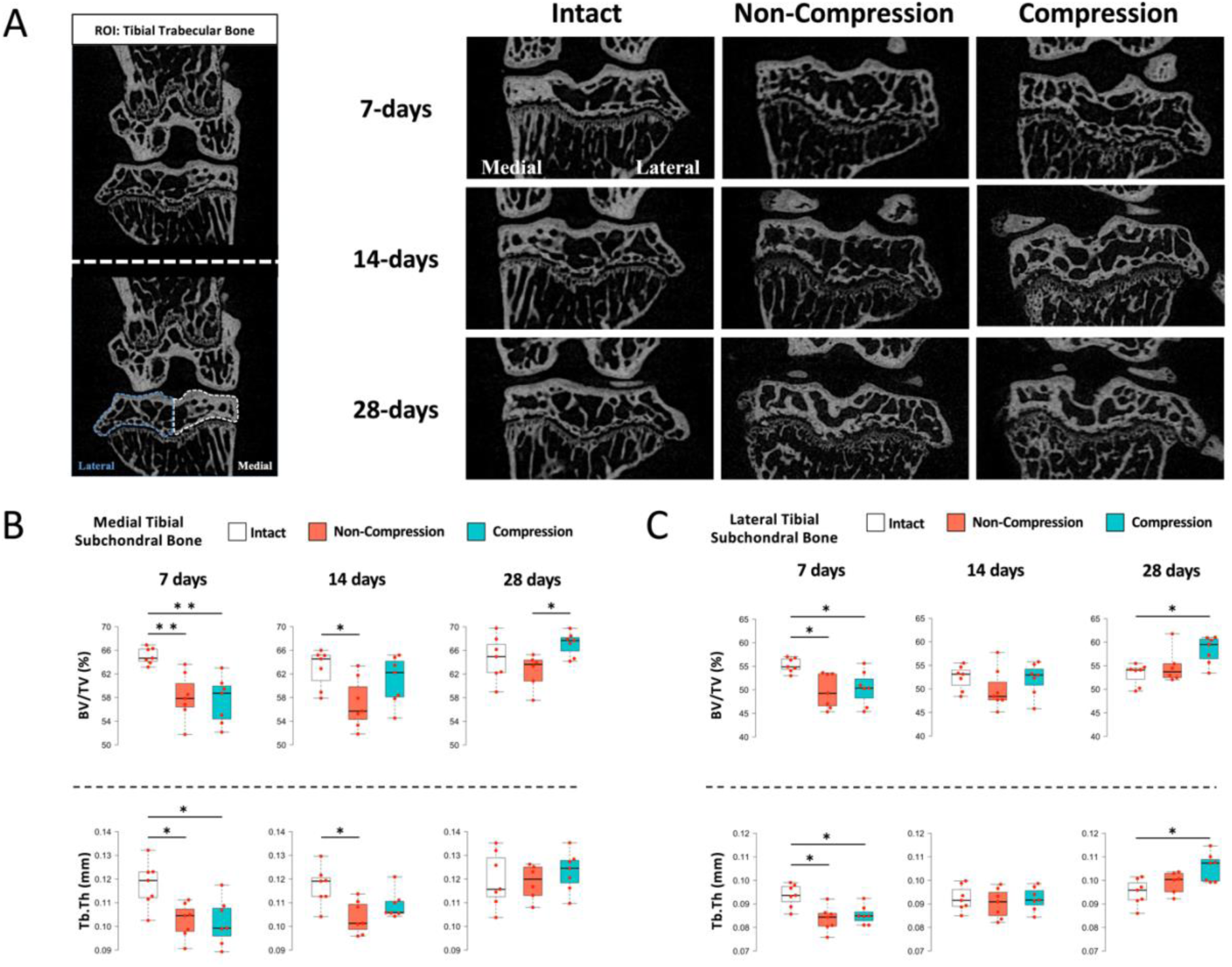
(A) The ROI of medial and lateral tibial trabecular bone and microCT images. (B) Analysis of subchondral bone microstructure in the medial and lateral tibial compartments. In the medial compartment, the Compression and the Non-Compression groups caused bone loss with a significant decrease in BV/TV and Tb.Th compared to the Intact group at 7 days. At 28 days, the Compression group showed significantly increased BV/TV compared to the Non-Compression group. In the lateral compartment, both ACL-R groups exhibited significant bone loss compared with the Intact group at 7 days. Unlike the medial compartment, the Compression group exhibited a significant increase of BV/TV and Tb.Th compared to the Intact group at 28 days. Data are presented as the median ± interquartile range. *P< 0.05. **P< 0.01.

### Histological Analysis for Cartilage Degeneration

Safranin-O/fast green staining was performed to evaluate articular cartilage degeneration in the medial and lateral tibial compartments using the Osteoarthritis Research Society International (OARSI) histopathological grading system ^25^. We initially assessed the whole joint pathology in the knee joint, then assessed the medial and lateral tibial plateau separately.

### Immunohistochemical Analysis

Immunohistochemical (IHC) staining was performed using anti-MMP-13 (1:200, bs-0575R, Bioss). Detailed protocols are described in the Supplementary Materials. We calculated the ratio between the number of MMP-13 positive cells and the number of chondrocytes in the anterior and posterior area of the articular cartilage with regions of interest of 40,000 µm^2^ (200 µm × 200 µm). As with the OARSI score, we initially evaluated the whole joint and then subsequently assessed the medial and lateral tibial plateau separately.

### Statistical Analysis

Statistical analysis was performed using RStudio. Student’s t-test was used for the osteophyte formation and Wilcoxon rank sum was used for the microscope and FRI data. One-way analysis of variance was performed for the joint laxity test and MMP-13 IHC analysis initially, and then the Tukey-Kramer test was used for post-hoc analysis. The Kruskal-Wallis test was used to compare subchondral bone µCT data and OARSI scores, and the Steel-Dwass method was used for the subsequent multiple comparisons. Parametric data are expressed as the mean ± 95% confidence intervals (95% CI); non-parametric data are expressed as the median ± interquartile ranges. Statistical significance was set at p < 0.05.

## RESULTS

### AP joint Laxity Test of the Knee

Compared with the Intact group, both the Compression and Non-Compression ACL-R groups had significantly increased AP joint laxity at both 60 ° and 90 ° (Figure 1B). However, no significant differences were observed between the Compression and Non-Compression groups at either angle. This suggests that any differences in OA progression between the ACL-R groups can be attributed to the initial effect of compressive force at the time of ACL injury.

### Microscope Analysis of Cartilage Roughness in the Tibial Plateau

Surface roughness in the region not covered by menisci was increased in the Compression ACL-R group, especially on the medial tibial plateau. The normalized arithmetical mean height of the medial compartment in the Compression group was significantly higher than that in the Non-Compression group, but no difference was observed in the lateral compartment (Figure 2A). In addition, there were no significant differences in the maximum height of the medial and lateral compartments between groups. These results indicate that compressive force during compression ACL injury caused initial micro-damage in the surface layer of articular cartilage.

### Quantification of MMP Fluorescence Intensity

At 3 days following injury, MMP-induced fluorescence intensity around ACL-R knees increased slightly in the Compression (Normalized data: 0.31 ± 0.659) and Non-Compression groups (0.512 ± 0.472) compared with each contralateral intact knee, and no significant difference was observed between groups (Figure 2B). Whereas, at 7 days, fluorescence intensity increased moderately in the Compression (0.684 ± 0.597) and slightly in the Non-Compression (0.17 ± 0.368) groups. However, there were no significant differences between these groups.

### MicroCT Measurement of Osteophyte Formation

Mineralized osteophyte volume was quantified at 14 and 28 days because no osteophytes were observed at 7 days following injury. Similarly, only a small amount of mineralized osteophyte formation was observed in both ACL-R groups at 14 days (Figure 3A). At 28 days following injury, a greater volume of ectopic mineralized tissue was observed in the medial femoral condyle and tibial plateau of injured joints. However, there were no significant differences between ACL-R groups.

### MicroCT Analysis of Epiphysis Trabecular Bone of the Distal Femur and Proximal Tibia

At 0 days post-injury, no micro injuries or fractures were observed in the femoral and tibial epiphysis trabecular bone in either ACL-R group, and there were no significant differences in the microstructure between any experimental groups (Supplementary Figure 2A-B). This indicates that compressive force while inducing ACL rupture didn’t cause concomitant structural failures in the subchondral trabecular bone. Femoral epiphysis trabecular bone showed a significant decrease of BMD, BV/TV, and Tb.Th in the Compression and Non-Compression groups compared with the Intact group at 7 days, which suggests acute bone absorption at an early time point in both ACL-R groups (Figure 3C). These parameters were still significantly lower in the Compression and Non-Compression groups at 14 days. However, the Compression group recovered bone volume moderately at 28 days, and no significant difference in the BMD was observed between the Intact group and the Compression group at this time point, although the Non-Compression group was still significantly lower than the Intact group. Tb.N and Tb.Sp results are described in the Supplementary Material (Supplementary Figure 3A).

Similarly, tibial epiphysis trabecular bone also exhibited a significant decrease in BMD and BV/TV in the Compression and Non-Compression groups at 7 days following injury (Figure 3D). At 14 days, the significantly decreased BMD and BV/TV was still present in the Non-Compression group, whereas the Compression group slightly increased bone volume and there was no significant difference in BV/TV between the Intact and the Compression groups. Furthermore, in the Compression group at 28 days, the bone volume increased to the same level as in the Intact group. The Non-Compression group still showed deficits in bone microstructure at this time point, therefore the BMD and BV/TV in the Intact and Compression group were significantly greater than in the Non-Compression group. Tb.N and Tb.Sp results are described in the Supplementary Material (Supplementary Figure 3B).

### MicroCT Analysis of Subchondral Bone in the Medial and Lateral Tibial Compartment

Analysis of subchondral bone microstructure in the medial and lateral tibial compartments at 7, 14, and 28 days (Figure 4A) showed that in the medial compartment, the Compression and the Non-Compression groups caused bone loss with a significant decrease in BV/TV and Tb.Th compared to the Intact group at 7 days (Figure 4B). Although BV/TV and Tb.Th in the Non-Compression group were still significantly lower than in the Intact group at 14 days, the Compression group increased BV/TV and Tb.Th and no differences were observed compared to other groups. Additionally, the Compression group exhibited significantly increased BV/TV compared to the Non-Compression group at 28 days.

In the lateral compartment, both ACL-R groups exhibited significant bone loss compared with the Intact group at 7 days similar to what was observed in the medial compartment (Figure 4C). At 14 days, the Compression and Non-Compression groups increased BV/TV and Tb.Th, and no differences were observed between any experimental groups. Unlike the medial compartment, the Compression group exhibited a significant increase of BV/TV and Tb.Th compared to the Intact group at 28 days. Tb.N and Tb.Sp results for both compartments are described in the Supplementary Materials (Supplementary Figure 4A-B).

### Histological Analysis of Cartilage Degeneration in the Medial and Lateral Tibial Compartment

Histological analysis did not detect any acute injuries of articular tissues other than the ACL (cartilage, menisci, subchondral bone) in the knee joint at 0 days following injury (Supplementary Figure 5). In the medial tibial plateau at 7 days, over half of the mice in the Compression group and some mice in the Non-Compression group developed a small loss of cartilage around the posterior edge, and cell proliferation and staining of Safranin-O were observed in the posterior region (Figure 5A). At 14 days, fibrillation of the cartilage surfaces, mild to moderate cartilage erosion, and osteophytes on the posterior tibia were observed in both ACL-R groups, especially in the posterior region. At 28 days, erosion extending to the center and posterior region of the subchondral bone and growth plate were observed in both ACL-R groups. These histological changes were observed in the lateral tibial plateau as well; however, the degree of OA progression was milder compared to the medial tibial plateau at all time points (Figure 5B).

**Figure 5.**
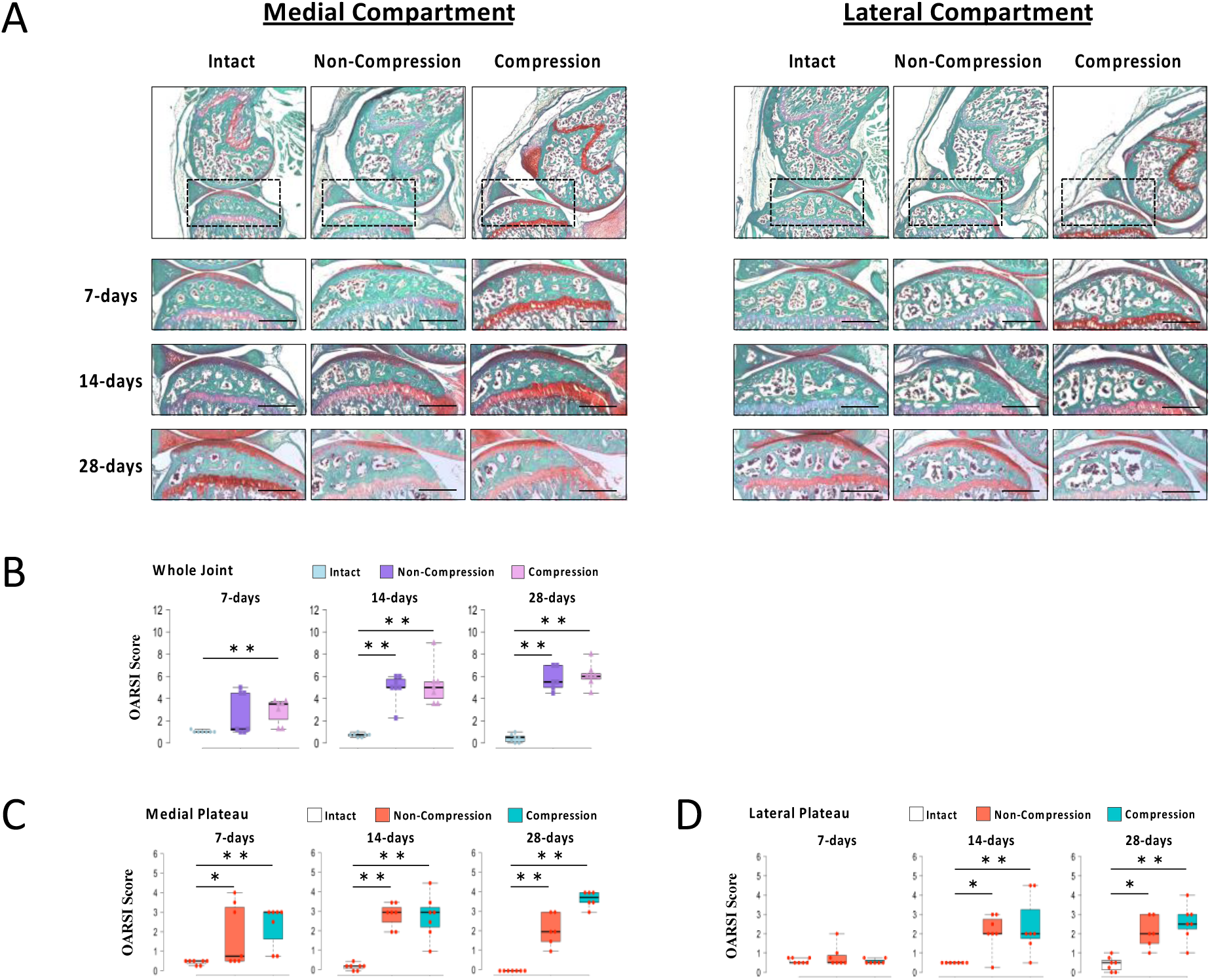
(A) Histological images of cartilage degeneration in the medial and lateral tibial compartment. (B) The OARSI score for the whole joint pathology in the Compression group was significantly higher than that in the Intact group at 7 days. At 14 and 28 days, both ACL-R groups significantly increased the OARSI score compared with the Intact group. (C) In the medial compartments, OARSI score in both ACL-R groups were significantly higher than that in the Intact group at all time points. (D) In the lateral compartment, both ACL-R groups caused a significant increase of OARSI score compared with the Intact group at 14 and 28 days. Data are presented as the median ± interquartile range. *P< 0.05. **P< 0.01. Black scale bar, 500 µm.

The OARSI score for the whole joint pathology in the Compression group increased significantly compared with the Intact group at 7 days (Figure 5C). Moreover, OARSI score in both ACL-R groups increased significantly compared with the Intact group at 14 days and 28 days. For the medial tibial plateau, OARSI score in both ACL-R groups were significantly higher than that in the Intact group at all time points (Figure 5D). For the lateral tibial plateau, OARSI score in both ACL-R groups were significantly higher than that in the Intact group at 14 days and 28 days (Figure 5E).

### Immunohistochemical Analysis of the Articular Cartilage

IHC analysis of MMP-13 was performed at 7 and 14 days following injury because cartilage was highly degenerated by 28 days (Figure 6A-B). At 7 days, the number of MMP-13-positive cells of the whole joint in the Compression group increased significantly compared with the Intact and Non-Compression groups (Figure 6C). Moreover, the Non-Compression group also significantly increased the positive cell ratio compared to the Intact group. Analysis of the medial tibial plateau showed similar results to the whole joint (Figure 6D), whereas for the lateral tibial plateau, the number of positive cells in the Compression group increased significantly compared with the other groups (Figure. 6E). At 14 days, the positive cells ratio of the whole joint in both ACL-R groups increased significantly compared with the Intact group. Similar results were observed in the medial and lateral tibial plateau.

**Figure 6.**
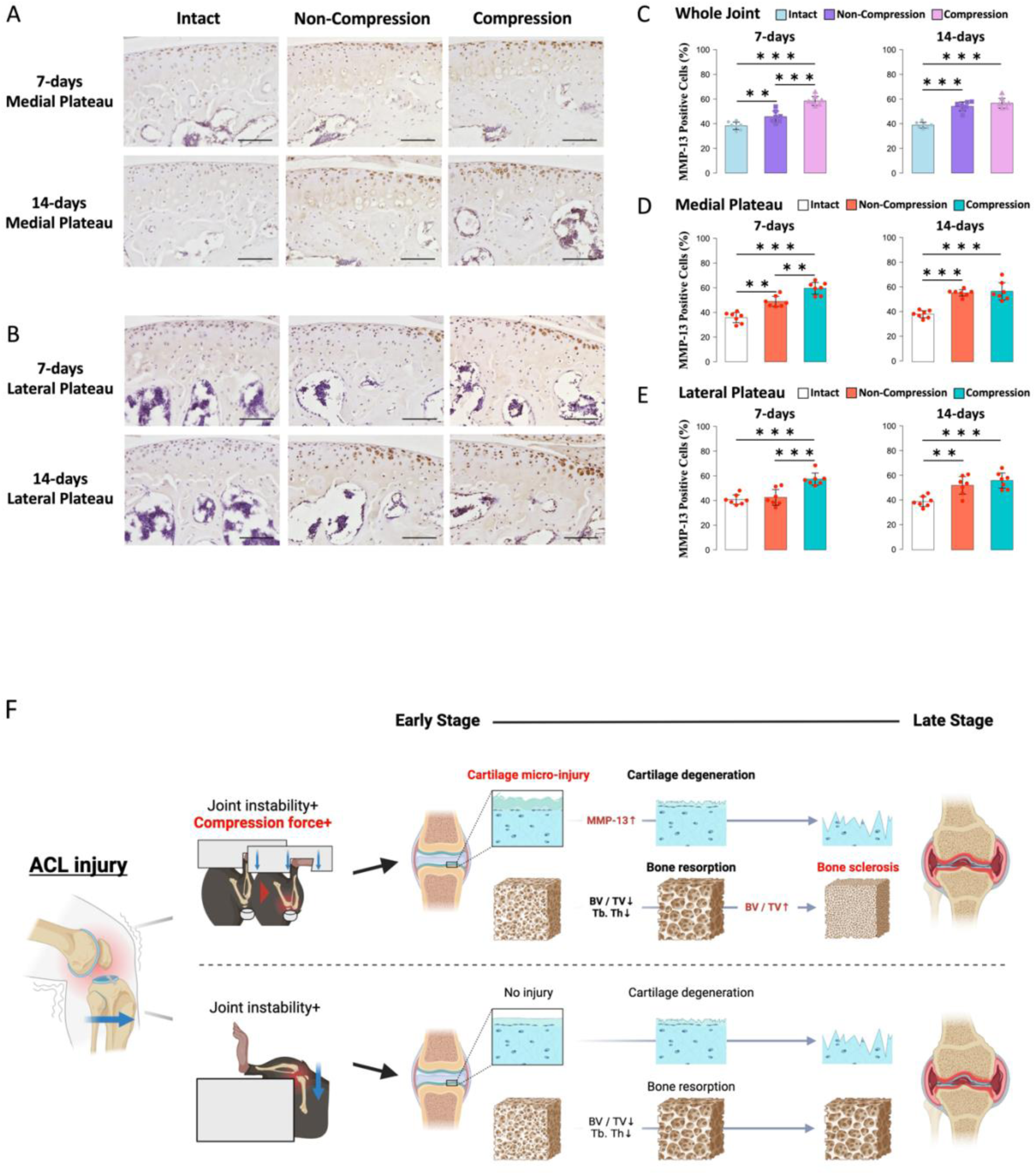
(A-B) Immunohistochemical images of MMP-13 in cartilage in the medial and lateral tibial plateau. (C-D) In the whole joint level and medial tibial plateau, Compression group increased the MMP-13 positive cell rate significantly compared with the Intact and Non-Compression groups at 7 days. Furthermore, the Non-Compression group also significantly increased the positive cell ratio compared to the Intact group. At 14 days, the positive cells ratio in both ACL-R groups increased significantly compared with the Intact group. (E) In the lateral tibial plateau at 7 days, the number of MMP-13 positive cells in the Compression group increased significantly compared with the other groups. At 14 days, both ACL-R groups increased the MMP-13 positive cell rate significantly compared with the Intact group. Data are presented as the mean ± 95% CI. **P< 0.01; ***P< 0.001. Black scale bar, 100 µm. (F) Schematic summarizing the mechanism of OA development following ACL injury with or without compression concomitant compression force. Compression-ACL rupture in mice caused initial micro-damage on the cartilage surface and early cartilage degeneration with MMP-13 production, leading to later bone formation in the subchondral bone.

## DISCUSSION

The mechanism of OA development following ACL injury involves two distinct processes: initial direct injury of the cartilage surface when the ACL is injured, and secondary joint instability due to the deficiency of the ACL. In this study, we compared the conventional Compression-induced ACL-R and Non-Compression ACL-R models, to clarify the individual effects of the initial injury response and secondary joint instability on OA progression after ACL injury. Although no significant differences in joint laxity were observed between ACL-R groups, the roughness of the cartilage after injury was increased significantly in the Compression ACL-R group but not the Non-Compression group. Similarly, MMP-induced fluorescence intensity at 7 days was higher in the Compression group than in the Non-Compression group. Moreover, the MMP-13 positive cell ratio in the Compression group was also increased significantly compared to the Non-Compression group at 7 days. Furthermore, sclerosis of tibial SCB in the Compression group developed significantly more than in the Non-Compression group at 28 days.

The ACL contributes to joint stability and ACL deficiency results in an increase in mechanical stress ^26^. Our assessment of AP joint laxity immediately after injury revealed that the Compression and Non-Compression groups had significantly increased joint laxity compared to the Intact group. However, no differences were observed between the ACL-R groups (Figure 1B). This result indicates that any differences in OA progression between the ACL-R groups can be attributed to the direct effect of compressive stress experienced while inducing ACL rupture.

Interestingly, microscopic analysis demonstrated that roughness on the surface of the medial tibial plateau in the Compression group was significantly increased compared to the Non-Compression group. However, no injuries such as cartilage erosion, meniscus injury, or subchondral bone loss were observed histologically and morphologically immediately following injury. A previous study using an intra-articular tibial plateau fracture mouse model reported that compression force to the tibial plateau to a target load of 55 N at a rate of 60 N/s caused massive soft tissue injuries, including to the meniscus, tendon, and cartilage ^27^. The mean compressive force applied with the Compression ACL-R model was approximately 10 N at a loading rate of 1 mm/s ^17^. The compressive force in the joint during Compression ACL-R was remarkably lower than in the previous study using the tibial plateau fracture model, therefore roughness on the cartilage surface indicated mild articular damage in the Compression group rather than severe damage.

In addition to the initial micro-injury, we also measured the biological response in the early stage using FRI with an MMP-activatable probe. MMP-3, MMP-9, and MMP-13 levels increased in animal models of osteoarthritis ^28–32^, and MMP-2, MMP-3, MMP-9, and MMP-13 increased in in vitro models of osteoarthritis ^33^, which can degrade collagens and proteoglycans of the articular cartilage. Compressive loading during ACL injury increases the production of matrix-degrading enzymes and inflammatory cytokines in chondrocytes, which results in chondrocyte apoptosis and cartilage degeneration ^5, 34, 35^. A previous animal study reported that even a single compressive load at 6 N without ACL rupture caused cartilage degeneration with chondrocyte apoptosis ^19^. However, in the current study no significant differences were observed between ACL-R groups; the Compression group showed a high MMP-induced fluorescence intensity using FRI at 7 days following injury. Furthermore, IHC analysis demonstrated that MMP-13 expression in chondrocytes of the Compression group increased significantly compared to the Non-Compression group. These molecular changes may be due to the initial micro-injury detected as surface roughness, which is consistent with early stage pathology of ACL injury in human patients. Interestingly, histological data also showed that the OARSI score in the whole joint of the Compression group significantly increased compared to the Intact group at 7 days. These results suggested that the compressive force on the cartilage surface while inducing ACL rupture may have accelerated cartilage degeneration in the early stage through the activation of MMPs. Alternatively, considering that there was no difference in OARSI score at 14 and 28 days, it is also possible that secondary joint instability is a more important factor in the progression of cartilage degeneration after the initial stage.

Subchondral bone provides mechanical and nutritional support for cartilage, and microenvironmental changes in subchondral bone might affect cartilage metabolism directly or indirectly ^36^. Generally, subchondral bone reacts to mechanical stress and is faster to remodel than articular cartilage due to its robust innervation and blood supply that provide it with a high capacity of turnover ^37, 38^. We hypothesized that subchondral bone would react to compressive force and induced structural changes during the early stage. Indeed, we observed loss of subchondral bone volume in both ACL-R groups following injury. However, the Compression group induced subchondral bone sclerosis in the medial tibial plateau compared to the Non-Compression group at 28 days, and the lesion area was consistent with the cartilage degeneration. Some reports have shown that cartilage degeneration precedes subchondral bone changes ^39–41^ and osteoarthritic chondrocytes enhanced osteoblast differentiation in subchondral bone via the ERK1/2 pathway ^42^. Our results revealed that ACL injury with concomitant compressive force initially caused micro-injury on the cartilage surface, then subchondral bone remodeling and sclerosis in the medial compartment at an earlier time point than ACL injury without compressive loading (Figure 6F). This earlier switch from bone loss to bone sclerosis is consistent with an accelerated PTOA progression in the Compression ACL-R group compared to the Non-Compression group.

There are two main limitations to this study. First, articular surface roughness analysis was only performed immediately after the induction of each model. Microscopic evaluation can determine the degree of cartilage degeneration as a surface rather than a line in more detail. Roughness analysis was able to detect micro-injury on the cartilage surface at 0 days (Figure 2A), which was not possible to see with histological observation (Supplementary Figure 5). Further roughness analysis at additional time points may enable us to reveal the reaction of cartilage degeneration to compression force, especially in the early phase. Second, we didn’t investigate the molecular biological mechanism of subchondral bone remodeling and sclerosis during PTOA progression. It has been shown that the proliferation and differentiation of osteoprogenitors are stimulated by platelet-derived growth factor-A, transforming gene factor-β1, and fibroblast growth factor-1, resulting in subchondral bone formation ^43^. We revealed that chondrocytes reacted to initial compression force and increased MMP-13; therefore, additional experiments about the biological mechanism should be performed to develop further understanding of these mechanisms.

In conclusion, concomitant joint contact while non-invasively inducing ACL rupture in mice caused initial micro-damage on the cartilage surface and early cartilage degeneration with MMP-13 production, leading to later bone formation in the subchondral bone. Understanding the initial pathology of the ACL injury may be an important indicator of disease etiology and represent a potential preventative approach for mitigating secondary OA development.

## Acknowledgments

The author(s) received no financial support for the research, authorship, and/or publication of this article. We would like to thank Editage (www.editage.com) for English language editing. The study design and summary scheme were created with BioRender.com. Research reported in this publication was partially supported by the National Institute of Arthritis and Musculoskeletal and Skin Diseases, part of the National Institutes of Health, under Award Number R01 AR075013.

## Author contributions

All authors approved the final submitted manuscript.

Study design: TK and BC

Making model and Data collection: KT and YL

Mechanical analysis: KT and BO

Fluorescent reflectance imaging: KT and YL

Morphological analysis: KT, YL, and KA

Histological analysis: KT, KA, and SE

Manuscript composition: KT, TK, and BC

