## supplemental materials for "Concurrent Joint Contact in Anterior Cruciate Ligament Injury induces cartilage micro-injury and subchondral bone sclerosis, resulting in knee osteoarthritis"

### Supplementary materials

#### Methods

##### Microscope analysis of cartilage surface roughness

For samples collected 0 days following injury, soft tissue was removed carefully to expose the tibial plateau, then 3D optical profilograph images of the whole tibial plateau were taken (VR-6000; Keyence, JPN). Intra-articular tissues such as ligament, menisci, synovium, and infrapatellar fat pad were removed carefully to expose the tibial plateau, and specimens were stored in the 1xPBS not to dry. The tibia was fixed on a table with clay so that the tibial diaphysis was vertical as much as possible, and the microscope images were taken in high-resolution mode. Initially, the reference lane was set by surrounding the joint surface, and the analysis areas of all samples were unified so that they were parallel (Supplementary Figure 1A-B). Then, we measured the surface roughness on the uncovered region by menisci with an ellipse of 0.6mm-1.2mm diameter because it was expected that compression force may induce micro-injury to the cartilage surface directly. The region of interest accounted for 38.5 ± 0.86% and 35.4 ± 0.69% in the medial and lateral tibial plateau, respectively. The arithmetical mean height and maximum height of the cartilage surface were measured by normalizing to the contralateral intact knee of each mouse.

A

B

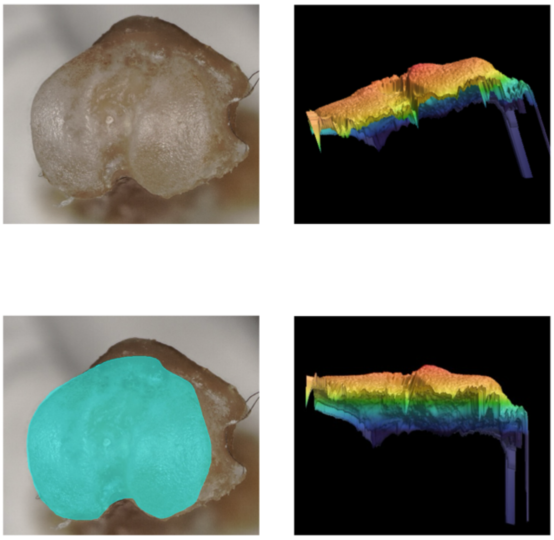

Supplementary Figure 1. (A) The tibial plateau fixed on a table with clay to be vertical, however, some samples were tilted. (B) After taking images, we set the reference lane by surrounding the joint surface, and the tibial plateau of all samples were unified to be parallel.

**Fluorescent reflectance imaging (FRI)**

At 3 and 7 days post-injury, mice were imaged in vivo using an optical imaging system (IVIS Spectrum, PerkinElmer, Waltham, MA). Based on the previous study, 10 mL (~0.1 mg/kg) of MMPSense was injected through retro-orbital sinus under anesthesia with isoflurane 24 h prior to imaging (1), and hair around the knee joint of both legs was removed with a depilatory. Mice were imaged in the knee extension position using the imaging system (IVIS Spectrum, PerkinElmer, Waltham, MA) and the ROI was set as a circle of 0.7 $mm^2$ that surrounded the tibial tuberosity to the superior border of the patella to exclude extra-articular soft tissue such as muscle. The legs were taped down across the ankle, and image processing and quantification were performed via IVIS Living Image software. The excitation and emission filters were set at 675/720 nm ex/em and the exposure time was 0.75 s. Spatial binning of pixels was set at the Medium option and the F/Stop was set to 2. Quantification of fluorescence intensity was performed by evaluating the total radiant efficiency ([photons/sec]/[mW/$cm^2$]) of the signal within a ROI. To unify the mouse-to-mouse variation in the delivery of the fluorescent probe, the radiant efficiency of the ACL-R knee was normalized to the contralateral intact knee of each mouse.

**Micro-Computed Tomography Analysis of Osteophyte Formation and Epiphyseal Bone Microstructure**

Bilateral knees were scanned using micro-computed tomography (µCT 35, SCANCO, Brüttisellen, Switzerland) with the following parameters: nominal voxel size = 10 µm, energy = 55 kVp, intensity = 72µA, integration time = 800 ms. For analysis of osteophyte formation at 7, 14, and 28 days after injury, VOI contouring included all heterotopic mineralized tissue around the joint, as well as the patella, fabellae, and menisci. The average bone volume of the patella, fabellae, and menisci in the Intact group was used as the "baseline" volume to calculate osteophyte formation in the Compression and Non-Compression groups. The difference in total bone volume between the injured mouse knees and the baseline volume was calculated to determine the total osteophyte volume for each injured joint.

We also assessed the morphological changes in epiphyseal trabecular bone at 0, 7, 14, and 28 days after injury. Morphological analysis of trabecular bone in the tibia and femur was performed by manually drawing contours on 2D transverse slices; the ROI was designed as the trabecular bone enclosed by the growth plate and subchondral cortical bone plate (Figure 3B). Additionally, whole subchondral bone including the cortical bone plate and trabecular bone in the medial and lateral tibial compartments were also measured (Figure 4A). Using the manufacturer's analysis software, we quantified apparent bone mineral density (BMD, $g/cm^3$), bone volume per total volume (BV/TV, %), trabecular number (Tb.N, 1/mm), trabecular thickness (Tb.Th, mm), and trabecular separation (Tb.Sp, mm).

**Immunohistochemical analysis**

To assess the expression of MMP-13, immunohistochemical staining was performed using the avidin-biotinylated enzyme complex method and a Vectastain Elite ABC Rabbit IgG Kit (Vector Laboratories, Burlingame, CA, USA). The tissue sections were deparaffinized with xylene and ethanol, and antigen activation was performed using proteinase K (Worthington Biochemical Co., Lakewood, NJ, USA) for 15 min. Endogenous peroxidase was inactivated with 0.3% H2O2/methanol for 30 min. Nonspecific binding of the primary antibody was blocked using normal goat serum for 30 min. The sections were then incubated with anti-MMP-13 antibody (1:200, bs-0575R, Bioss) overnight at 4C. Afterward, the sections were incubated with biotinylated secondary anti-rabbit IgG antibody and stained using Dako Liquid DAB þ Substrate Chromogen System (Dako, Glostrup, Denmark). The cell nuclei were stained using hematoxylin at a concentration of 25%. We calculated the ratio between the number of MMP-13 positive cells and the number of chondrocytes in the anterior and posterior area of the articular cartilage with regions of interest of 40,000 $\mu m^2$ (200 mm × 200 mm). We initially evaluated the whole joint and then subsequently assessed the medial and lateral tibial plateau separately.

### Results

##### µCT measurement of osteophyte formation

We evaluated the concomitant injuries by compression force at 0 days post-injury, however, no micro injuries or fractures were observed in the femoral and tibial epiphysis trabecular bone in all groups. Moreover, there were no significant differences in the BMD, BV/TV, Tb.N, Tb.Th, and Tb.Sp. between all groups (Supplementary Figure 2A-B).

A

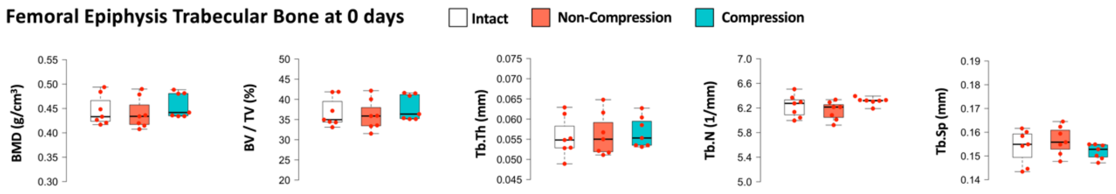


B

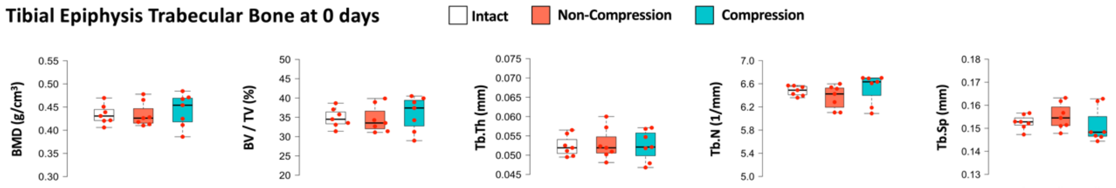


Supplementary Figure 2. (A-B) No significant differences in the BMD, BV/TV, Tb.N, Tb.Th, and Tb.Sp were observed in the femoral and tibial epiphysis trabecular bone at 0 days. Data are presented as the median ± interquartile range.

**μCT analysis of epiphysis trabecular bone of the distal femur and proximal tibia**

Femoral epiphysis trabecular bone showed that both ACL-R groups decreased Tb.N and increased Tb.Sp significantly compared with the Intact group at 7 days and 14 days (Supplementary Figure 3A). At 28 days, Tb.N in the Compression and Non-Compression groups was still significantly lower than that in the Intact group, whereas Tb.Sp in the Compression group decreased and that in the Non-Compression group was significantly higher compared with the Intact group. Tibial epiphysis trabecular bone showed no significant differences in the Tb.N and Tb.Sp between all groups at 7 days, however, Tb. N in the Non-Compression group decreased significantly compared to the Intact group at 14 and

28 days (Supplementary Figure 3B). Tb.Sp increased significantly in the Non-Compression group compared to the Intact and Compression group at 28 days.

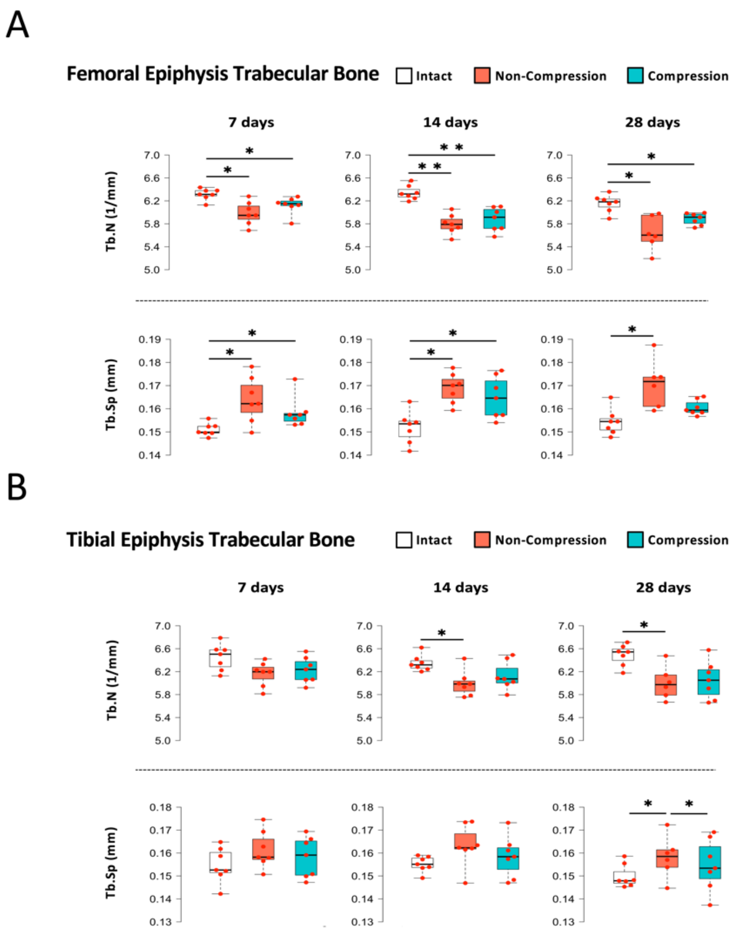


Supplementary Figure 3. (A) In the femoral epiphysis trabecular bone, both Compression and Non-Compression ACL-R groups decreased Tb.N and increased Tb.Sp significantly compared with the Intact group at 7 days and 14 days. Tb. N in the both ACL-R groups still significantly lower at 28 days. Although Tb. Sp in the Non-Compression group still significantly increased compared with the Intact group, no differences were observed between the Compression and Intact groups. (B) In the tibial epiphysis trabecular bone, Tb. N in the Non-Compression group decreased significantly compared with the Intact group at 14 and 28 days. Tb.Sp in the Non-Compression group was significantly higher than that in the Intact and Compression groups at 28 days. Data are presented as the median ± interquartile range. *P< 0.05. **P< 0.01.

In the tibial subchondral bone, there were no significant differences in Tb.N and Tb. Sp of the medial and lateral compartments between all groups at all time points (Supplementary Figure 4A-B).

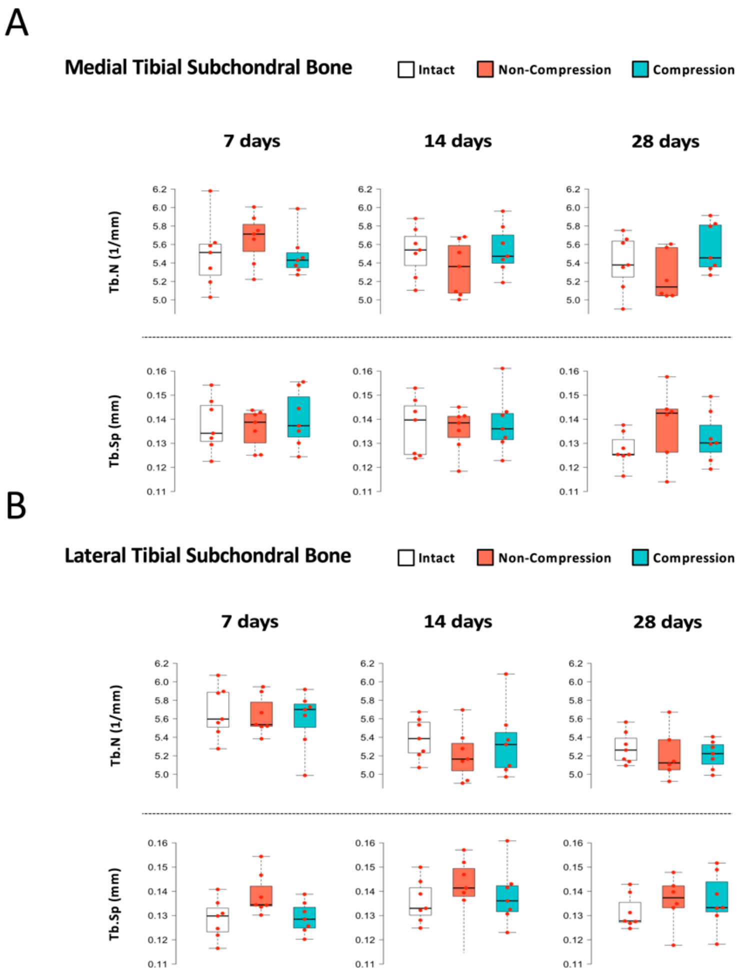

Supplementary Figure 4. (A-B) In the medial and lateral tibial subchondral bone, no significant differences in Tb.N and Tb. Sp were observed between all groups at all time points. Data are presented as the median ± interquartile range.

**Histological analysis of cartilage degeneration in the medial and lateral tibial compartment**

We performed histological observation in the medial and lateral compartments at 0 days immediately after making the Non-Compression and Compression models. In all mice, no histological injuries of intra-articular tissues, such as cartilage, menisci, and SCB were confirmed in the whole knee joint (Supplementary Figure 5).

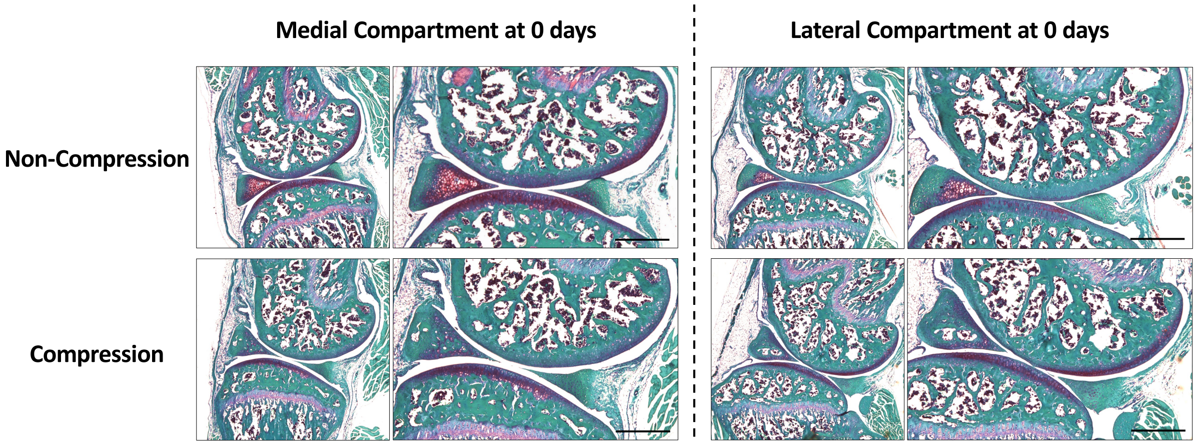


Supplementary Figure 5. Compression and Non-Compression ACL-R groups induced no histological articular damages in the whole knee joint at 0 days following injury. Black scale bar, 500μm.
